# Role of gut epithelial-cell derived transglutaminase 2 in the formation of celiac disease autoantibodies

**DOI:** 10.64898/2026.09.01.748464

**Authors:** Runa I. Løberg, Helena A. Abdi-Dezfuli, Liv Kleppa, Alisa E. Dewan, Maureen T. Meling, Ludvig M. Sollid, M. Fleur du Pré

## Abstract

Formation of autoantibodies to transglutaminase 2 (TG2) in celiac disease likely involves TG2-gluten complexes that allow gluten-specific CD4^+^ T cells to provide help to TG2-specific B cells. To investigate whether TG2 derived from intestinal epithelial cells (IECs) contributes to this process, we generated mice with inducible IEC-specific expression of TG2 fused to a deamidated gluten peptide (DGP) containing a T-cell epitope. Upon induction, the TG2-DGP fusion protein was expressed in IECs and released into the intestinal lumen. In HLA-DQ2.5 transgenic mice with activated gluten-specific CD4^+^ T cells and naive TG2-specific B cells, expression of TG2-DGP was immunogenic and drove the production of intestinal and systemic anti-TG2 autoantibodies. TG2-specific B cells predominantly expanded in Peyer’s patches, suggesting that their priming and subsequent differentiation into lamina propria TG2-specific IgA^+^ plasma cells occurs in gut-associated lymphoid tissues. The findings demonstrate immunogenicity of IEC-derived TG2-DGP fusion antigen supporting the notion of a pathogenic role for luminal TG2 in driving anti-TG2 autoimmunity in celiac disease.

**IMPACT STATEMENT:** Luminal release of intestinal epithelial cell-derived transglutaminase 2–gluten fusion protein initiates anti-TG2 autoimmunity by linking gluten-specific T-cell help to TG2-specific B cells in Peyer’s patches.

## INTRODUCTION

Celiac disease is a prevalent immune-mediated disorder driven by ingestion of gluten proteins that is hallmarked by autoantibodies specific for transglutaminase 2 (TG2) (Dieterich et al., 1997). The lesion in the small intestine is characterized by villous blunting and crypt cell hyperplasia with increased epithelial cell turnover and infiltration of inflammatory cells both in the epithelium and in the lamina propria. The plasmacytosis in the lamina propria is particularly striking with 10-20% of the plasma cells being specific for TG2 and 0.5-1.0% being specific for deamidated gluten peptides (DGPs) (Lindeman et al., 2021; Steinsbo et al., 2014).

TG2 is a calcium dependent enzyme that modifies glutamine residues in peptides in a sequence specific manner. It forms a transient enzyme-substrate intermediate to either deamidate the glutamine residue to a negatively charged glutamate or to crosslink it to a primary amine, which can be a lysine residue of another peptide, resulting in transamidation and isopeptide formation. Many gluten peptides are excellent substrates for TG2 and are readily deamidated by TG2 (Fleckenstein et al., 2004). Such deamidated gluten peptides are recognized by CD4^+^ T cells of celiac disease patients when presented by the celiac disease-associated human HLA molecules HLA-DQ2.5, HLA-DQ2.2 or HLA-DQ8 (Iversen and Sollid, 2023). These HLA-DQ allotypes are required for celiac disease to develop, and they all have a preference for binding negatively charged peptide ligands, explaining the HLA association of the disease. The gluten-specific CD4^+^ T cells, when activated in the gut mucosa, instigate immune reactions that eventually lead to formation of the disease lesion. Strikingly, the formation of antibodies to TG2 only occurs in individuals who carry the celiac disease-associated HLA-DQ allotypes and who consume gluten. The dependence on HLA and gluten for the autoantibody production can be explained by involvement of enzyme-substrate TG2-gluten complexes. Such complexes can act in a hapten-carrier like manner, with TG2 as the hapten and gluten as the carrier, thereby allowing TG2-specific B cells to interact with and receive T-cell help from gluten-specific CD4^+^ T cells (Sollid et al., 1997). While both TG2-gluten enzyme-substrate intermediate complexes and gluten peptides crosslinked to TG2 via isopeptide bonds are likely able to activate gluten-specific T cells, accumulating evidence suggests that the enzyme-substrate complex is the most relevant form in the pathogenesis of celiac disease (du Pre et al., 2024). Of note, the amount of enzyme-substrate complex formed is dependent on the substrate concentration, with high substrate concentration giving high occupancy of the enzyme’s active site.

Despite this detailed insight into the formation of gluten T-cell epitopes and autoantibodies to TG2, it remains unknown where and how the complexes of TG2 and gluten are generated. TG2 is ubiquitously expressed and is located mainly intracellularly in the cytosol (Lorand and Graham, 2003). While there is literature suggesting that TG2 is found extracellularly and is involved in extracellular matrix formation (Belkin, 2011), we observed no signs of B-cell tolerance in Ig knock-in (KI) mice that express a celiac patient-derived anti-TG2 B-cell receptor (BCR), suggesting that developing and circulating B cells are minimally exposed to extracellular TG2 under normal conditions (du Pre et al., 2020). To account for this observation and to place TG2 at a location where there is a high concentration of gluten peptides, we proposed that the pathogenic TG2 in celiac disease derives from shed intestinal epithelial cells (IECs) (Iversen et al., 2020). A large number (approximately 2 x 10^10^) of IECs are shed from the tip of the villi into the intestinal lumen each day as part of the continuous renewal of the epithelial layer (Sender and Milo, 2021). Most shed IECs are viable (Bahar Halpern et al., 2023; Glaeser et al., 2002; Meling et al., 2024), and can release their cytosolic contents upon disintegration. We have previously shown that both mouse and human IECs express fairly high levels of TG2 (Amundsen et al., 2023; Iversen et al., 2020), and that TG2 in IEC lysates can facilitate the interaction of TG2-specific B cells with gluten-specific CD4^+^ T cells when incubated with a substrate gluten peptide (Iversen et al., 2020). Moreover, IEC-derived TG2 present in the gut lumen of mice was demonstrated to be catalytically active and capable of deamidating gluten peptides in the presence of protease inhibitors, indicating that transient TG2-gluten complexes can be formed in the gut lumen, yet it remained unproven whether TG2 released from IECs *in vivo* stays intact for recognition by conformationally sensitive B-cell receptors (Meling et al., 2024). The model suggests that luminal TG2-gluten complexes are taken up by immune cells in gut-associated lymphoid tissue (GALT) in the same way as other luminal antigens derived from food or the microbiota, and in support of this notion it was demonstrated that TG2-specific B cells in the Peyer’s patch subepithelial dome area can sample TG2 that was added into ligated intestinal loops (du Pre et al., 2025).

To address whether IEC-derived TG2 can induce anti-TG2 autoimmunity *in vivo*, we generated mice with inducible expression of a TG2-DGP fusion protein in IECs. Upon induction, the TG2-DGP fusion protein is released into the intestinal lumen from shed IECs, providing a source of luminal TG2 antigen. In the presence of inflammatory gluten-specific CD4^+^ T cell help, luminal TG2-DGP is strongly immunogenic and initiates a robust anti-TG2 autoantibody response. These findings provide mechanistic support for a model in which IEC-derived luminal TG2 is pathogenic in celiac disease, driving the collaboration between TG2-specific B cells and gluten-specific CD4^+^ T cells through the formation of TG2-gluten complexes in the gut lumen.

## RESULTS

### Generation of mice with inducible expression of TG2-ω18merEE in IECs

To experimentally address the role of TG2 derived from shed IECs in celiac disease, we generated a mouse strain with doxycycline (dox)-inducible (Tet-On) expression of a TG2-DGP fusion protein in IECs. The inducible transgene encoded an 18mer deamidated ω-gliadin peptide QPEQPF<u>PQPEQPFPW</u>QPQ harboring the immunodominant DQ2.5-glia-ω2 T-cell epitope (9mer core sequence underlined) which was fused to the C-terminus of TG2 via a flexible glycine linker (hereafter referred to as TG2-ω18merEE). The gene sequence encoding this TG2-ω18merEE fusion protein was connected to a nuclear histone H2B- eGFP fusion reporter gene via 2A peptide sequences and cloned into a targeting vector under the tetracycline responsive element (TRE)-tight promoter. This gene construct (Figure 1A) was subsequently integrated into the ROSA26 locus by gene targeting in C57Bl/6 embryonic stem cells, resulting in the generation of ROSA26-*Tgm2*-ω18merEE and H2B-eGFP transgenic mice (referred to as *Tgm2*-ω18merEE single transgenic (STG) mice). These mice were then crossed to Villin1-reverse tetracycline controlled transactivator (rtTA)*M2^V5^ transgenic mice (Chen et al., 2014), selectively expressing rtTA in IECs, (referred to as IEC-rtTA STG mice) to generate *Tgm2*-ω18merEE IEC-rtTA double transgenic (DTG) mice (referred to as DTG mice) (Figure 1B). In DTG mice, treatment with dox, a tetracycline analogue that activates the Tet-On system by binding to rtTA, induces expression of the TG2-ω18merEE fusion protein and the eGFP reporter specifically in the intestinal epithelium. Consistently, dox treatment led to robust expression of TG2-ω18merEE and eGFP in IECs (Figure 1C).

**Figure 1.**
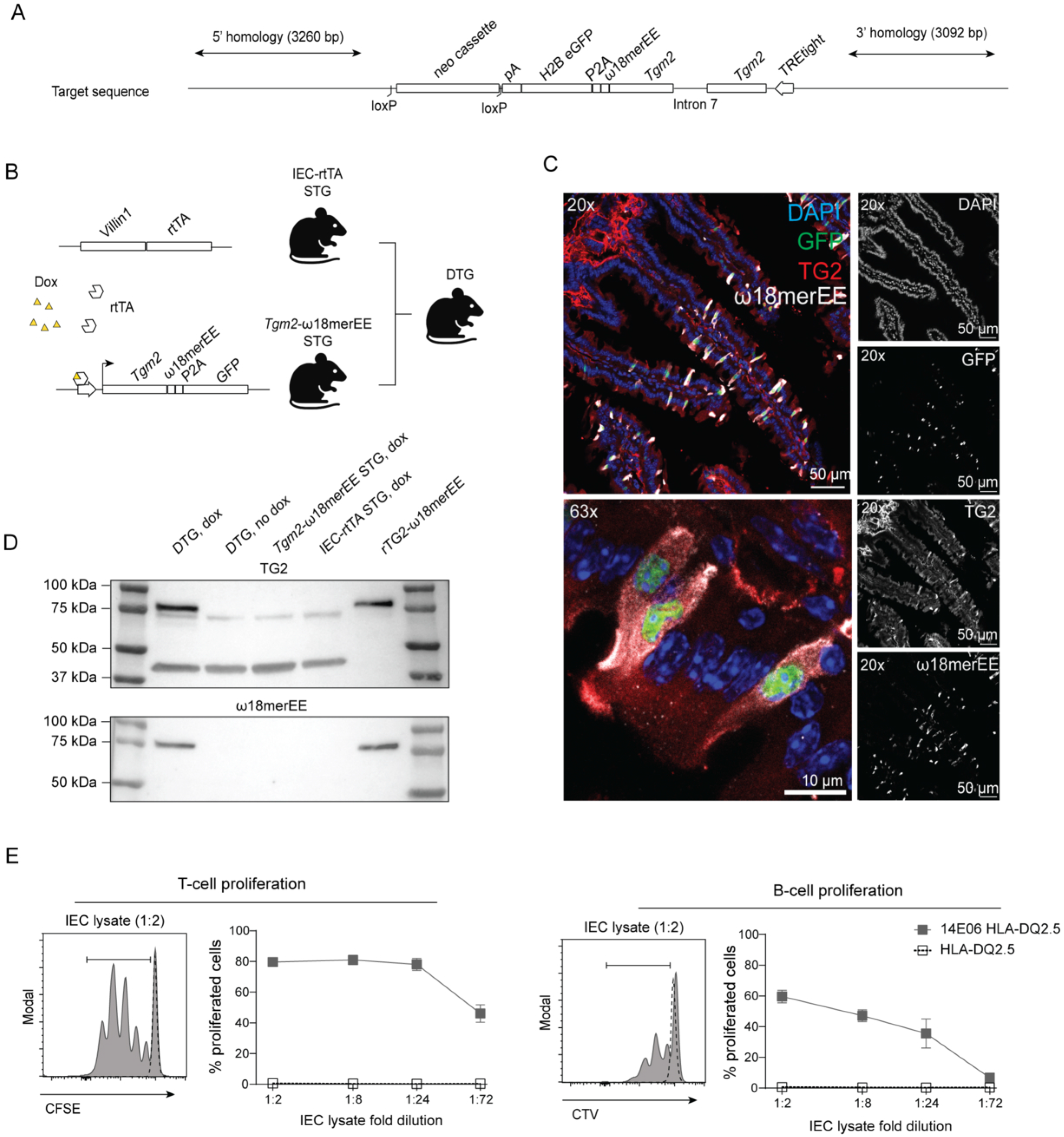
TG2-ω18merEE fusion protein is expressed in intestinal epithelial cells in doxycycline-treated double transgenic mice. **(A)** Schematic overview of the transgene construct made to generate ROSA26-*Tgm2*-ω18merEE; H2B-eGFP transgenic mice, referred to as *Tgm2*- ω18merEE single transgenic (STG) mice. The cDNA sequence encoding murine transglutaminase 2 (TG2), with retention of intron 7, was fused to cDNA encoding the 18mer deamidated gluten peptide QPEQPFPQPEQPFPWQPQ (ω18merEE) and linked to a nuclear histone H2B-eGFP reporter via P2A peptide sequences, all placed under control of the tetracycline-responsive promoter (TREtight) and targeted in reverse orientation into the *Rosa26* locus. **(B)** Illustration of doxycycline (dox)-induced expression of the transgenic TG2-ω18merEE fusion protein in intestinal epithelial cells (IECs) using the Tet-On system. In IEC-rtTA STG mice, expression of the reverse tetracycline-controlled transactivator (rtTA) is driven by the *Villin1* promoter, active in IECs. Crossing IEC-rtTA STG mice with *Tgm2*- ω18merEE STG mice generates double transgenic (DTG) mice, in which rtTA binds and activates the TREtight promoter in the presence of dox, thereby inducing TG2- ω18merEE expression in IECs. **(C)** Induction of eGFP and TG2-ω18merEE protein in small intestinal tissue from DTG mice 24 h after intraperitoneal administration of dox. eGFP (green) was detected by direct epifluorescence, and tissue was stained with anti-TG2 (red) anti-DGP (white) and DAPI (blue). **(D, E)** DTG, *Tgm2*-ω18merEE STG or IEC-rtTA STG mice were administered dox via drinking water (2 mg/ml) or normal drinking water (no dox) for 3 days followed by 24 h on normal drinking water. IECs were isolated from the small intestine and lysed by three freeze-thaw cycles. **(D)** Western blot detection of TG2-ω18merEE fusion protein in IEC lysates. Top: Anti-TG2 antibodies detect transgenic TG2-ω18merEE protein (∼82 kDa, upper band) and endogenous TG2 (∼78 kDa, lower band). The band at ∼42 kDa represents β-actin, which was included as loading control. Bottom: Anti-deamidated ω18mer (ω18merEE) antibodies detect the ω18merEE peptide in the transgenic TG2-ω18merEE fusion protein. Recombinant TG2-ω18merEE protein was loaded as a control. Data are representative of three independent experiments. **(E)** TG2-specific B cells and non-cognate B cells were isolated from *Tgm2*^-/-^ HLA-DQ2.5 14E06 mice and HLA-DQ2.5 mice, respectively, labeled with CellTrace Violet (CTV), and incubated with IEC lysate from dox-treated DTG mice for 10 minutes on ice. B cells were washed and co-cultured with CFSE-labeled gluten-specific CD4^+^ T cells, isolated from HLA-DQ2.5 TCR-glia-ω2 mice, for 3 days at 37°C. Proliferation of T cells (left) and B cells (right) as indicated by dilution of proliferation-tracking dyes is shown as the mean of culture duplicates ± SD. Data are representative of three independent experiments. **Source Data 1.** Uncropped Western Blots shown in Figure 1D. **Figure Supplement 1.** Gating strategy for Figure 1E.

To determine whether the IEC-expressed TG2-ω18merEE fusion protein can facilitate cognate interactions between TG2-specific B cells and gluten-specific CD4^+^ T cells, we performed an *in vitro* proliferation assay using IEC lysates of dox-treated DTG mice. Western blot analysis using antibodies against both TG2 and DGP confirmed the presence of the TG2-ω18merEE fusion in IEC lysates of dox-treated DTG mice, whereas no fusion protein was detected in IEC lysates from untreated DTG mice or from dox-treated STG control mice (Figure 1D). TG2-specific B cells were isolated from *Tgm2*^-/-^ HLA-DQ2.5 14E06 Ig KI mice that express a celiac-patient derived anti-TG2 BCR (du Pre et al., 2020), and gluten-specific CD4^+^ T cells were obtained from gluten-specific TCR transgenic mice expressing the immunodominant DQ2.5-glia-ω2-specific TCR (Lindstad et al., 2021). As we have previously shown that cytosolic TG2 is readily released during tissue disruption and B-cell isolation, leading to artifactual binding of TG2 by the BCR on freshly-isolated TG2-specific B cells (du Pre et al., 2020), we used HLA-DQ2.5 14E06 KI mice crossed with *Tgm2*^-/-^ deficient mice (De Laurenzi and Melino, 2001) as B-cell donors in all our experiments. TG2-specific B cells and gluten-specific CD4^+^ T cells were incubated with serial dilutions of IEC lysate from a dox-treated DTG mouse. Strong proliferative responses of both CD4^+^ T cells and B cells were observed, indicating that TG2-specific B cells efficiently took up the TG2-ω18merEE fusion protein and presented the DGP to the T cells and that the gluten-specific CD4^+^ T cells in turn provided T-cell help and induced proliferation of TG2-specific B cells (Figure 1E). No proliferation was observed when B cells isolated from regular HLA-DQ2.5 KI mice were used, indicating that uptake of the TG2-ω18merEE fusion protein was dependent on the TG2-specific BCR (Figure 1E). Collectively, these data demonstrate that functional TG2-ω18merEE fusion protein is expressed in an inducible manner in IECs of DTG mice upon dox administration.

### IEC-derived TG2-ω18merEE is detected in the gut lumen

Using a conditional KO mouse that specifically lacks TG2 expression in IECs, we have shown that TG2 present in the gut lumen in mice is derived from shed IECs (Meling et al., 2024). We therefore asked whether the transgenic TG2-ω18merEE fusion protein could be detected in the intestinal lumen. To determine the kinetics of transgenic protein expression, we assessed the presence of TG2-ω18merEE over time following a single i.p. injection of dox. The TG2-ω18merEE fusion protein was efficiently recovered from small-intestinal lavage fluids through immunoprecipitation using an antibody recognizing the gluten-epitope sequence of the fusion protein followed by Western blot protein visualization using an anti-TG2 antibody. The presence of TG2-ω18merEE fusion protein in the intestinal lumen could be detected from 12 h after dox administration, with levels becoming undetectable by day 5 (Figure 2).

**Figure 2.**
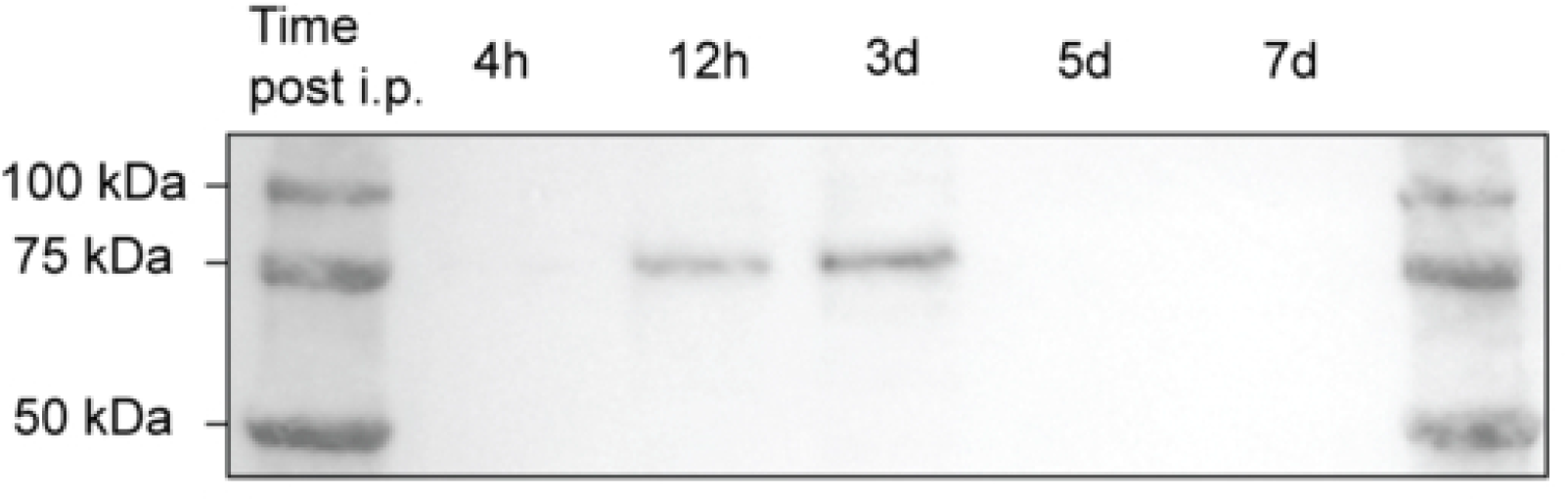
TG2-ω18merEE is detected in the intestinal lumen of dox-treated DTG mice. DTG mice received dox (50 mg/kg) by i.p. injection, and were euthanized 4 h, 12 h, or 3, 5 or 7 days after treatment. Small-intestinal lavage fluids were collected, and the TG2-ω18merEE fusion protein was extracted by immunoprecipitation using an antibody recognizing the deamidated ω-gliadin peptide, followed by Western blot visualization using a polyclonal anti-TG2 antibody. Data are representative of two independent experiments. **Source Data 2.** Uncropped Western Blot.

### IEC-derived TG2-ω18merEE induces anti-TG2 autoimmunity

We next assessed whether the IEC-derived TG2-ω18merEE fusion protein is sufficiently immunogenic to initiate an anti-TG2 autoantibody response. To this end, we developed an adoptive transfer model using DTG mice crossed to HLA-DQ2.5 KI mice as recipients. CD4^+^ gluten-specific T cells isolated from HLA-DQ2.5 KI TCR-glia-ω2 transgenic mice were first transferred and primed by intragastric (i.g.) immunization with DGP and cholera toxin (CT) as adjuvant (Figure 3A). Three days later, TG2-specific B cells isolated from *Tgm2*^-/-^ HLA-DQ2.5 14E06 KI mice were transferred, and TG2-ω18merEE expression was induced by i.p. dox injection. Dox-treated mice developed serum TG2-specific IgG and IgA that were detectable 7 days after dox administration (Figure 3B). TG2-specific IgA was also detected in small-intestinal lavage fluids, indicating a strong mucosal autoantibody response (Figure 3B). No anti-TG2 antibodies were detected in mice not administered dox, or in dox-treated *Tgm2*-ω18merEE STG mice (Figure 3B). The antibody response was strictly dependent on prior activation of the gluten-specific T cells, as DTG mice that received gluten-specific CD4^+^ T cells but were not immunized with DGP / CT did not develop anti-TG2 autoantibodies (Figure 3B). Consistent with the presence of autoantibodies in small-intestinal lavage fluids, TG2-specific plasma cells were detected in the small-intestinal lamina propria Figure 3C). Thus, expression of a TG2-ω18merEE fusion protein in IECs gives rise, in the presence of activated gluten-specific CD4^+^ T cells, to an anti-TG2 autoantibody response.

**Figure 3.**
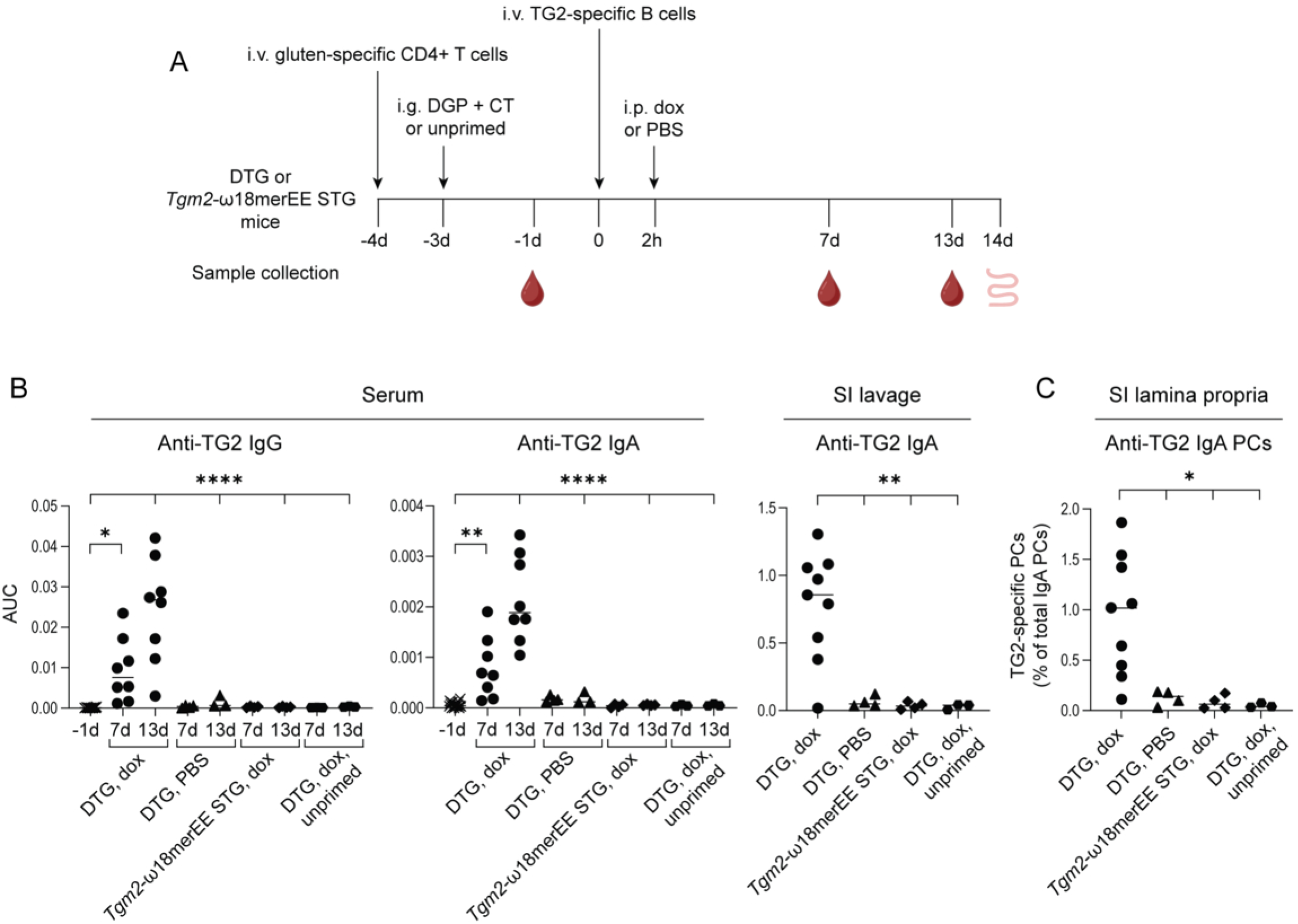
IEC-derived TG2-ω18merEE fusion protein drives anti-TG2 responses in a T-cell dependent manner. **(A)** Schematic overview of the experiment. Naive gluten-specific CD4^+^ T cells, isolated from HLA-DQ2.5 TCR-glia-ω2 mice, were transferred to HLA-DQ2.5 DTG or HLA-DQ2.5 *Tgm2*-ω18merEE STG recipient mice. The following day, T cells were activated by intragastric (i.g.) administration of deamidated gluten peptides (DGP, 100 µg or 500 µg) and cholera toxin (CT, 10 µg). Three days later, naive TG2-specific B cells isolated from *Tgm2*^-/-^ HLA-DQ2.5 14E06 KI mice were transferred and dox (50 mg/kg) was administered i.p. 2 hours after B-cell transfer. Serum was collected at the indicated time points. The small intestine was harvested 14 days (14d) after dox injection for collection of lavage fluids and isolation of lamina propria lymphocytes. **(B)** Anti-TG2 IgG and IgA in serum, and anti-TG2 IgA in small-intestinal (SI) lavage in DTG mice treated with dox or PBS, or in *Tgm2*-ω18merEE STG mice. The control group represents dox-treated DTG mice that did not receive i.g. immunizations of DGP and CT after T-cell transfer (unprimed). Area under the curve (AUC) was determined as a function of serum-to-buffer ratio measured as peaks above baseline, ignoring peaks < 10% of maximum. **(C)** Anti-TG2 IgA-producing plasma cells (PCs) expressed as a percentage of total IgA-producing PCs in the SI lamina propria in the groups as described under (**B**). (**B-C**) Data are shown with median and are pooled from five independent experiments. *p < 0.05, **p < 0.01, and ****p < 0.0001 by one-way ANOVA.

### Activated TG2-specific B cells are primarily found in Peyer’s patches

We next analyzed antigen-specific B-cell and T-cell responses at gut-associated immune inductive sites. CD45.1/CD45.2 congenic markers were used to track transferred B and T cells. In mice with dox-induced IEC expression of the TG2-ω18merEE fusion protein, TG2-specific B cells expanded in both Peyer’s patches (PPs) and gut-draining mesenteric lymph nodes (MLNs), but expansion of TG2-specific B cells was markedly stronger in PPs than in MLNs 7 days after dox administration (Figure 4A,B). In PPs, the vast majority of TG2-specific B cells were IgD^-^ (>90%) (Figure 4C), consistent with antigen-driven activation, and had class-switched to both IgG1 and IgA, with IgG1 cells dominating (Figure 4D). In MLNs of dox-treated DTG mice approximately 80% of TG2-specific B cells were IgD^-^ (Figure Supplement 2A). Only a very small fraction had switched to IgA indicating that IgA class switching is largely confined to PPs (Figure Supplement 2B). Notably, TG2-specific B cells in PPs displayed a germinal center phenotype (B220^+^ CD38^-^ CD95^+^ Bcl-6^+^ Ki-67^+^) (Figure 4E), and cells with this phenotype were also detected in MLNs (Figure Supplement 2C). Only very small proportions of transferred TG2-specific cells were detected in control groups (Figure 4A,B), and these cells mostly retained a naive phenotype (Figure 4C, Figure Supplement 2A). Taken together, these data suggest that PPs rather than MLNs are the primary inductive site for TG2-specific B-cell activation and class switching in response to IEC-derived TG2-ω18merEE. Of note, TG2-specific B cells in MLNs may reflect cells that were initially primed in PPs and subsequently migrated to the downstream gut-draining lymph nodes.

**Figure 4.**
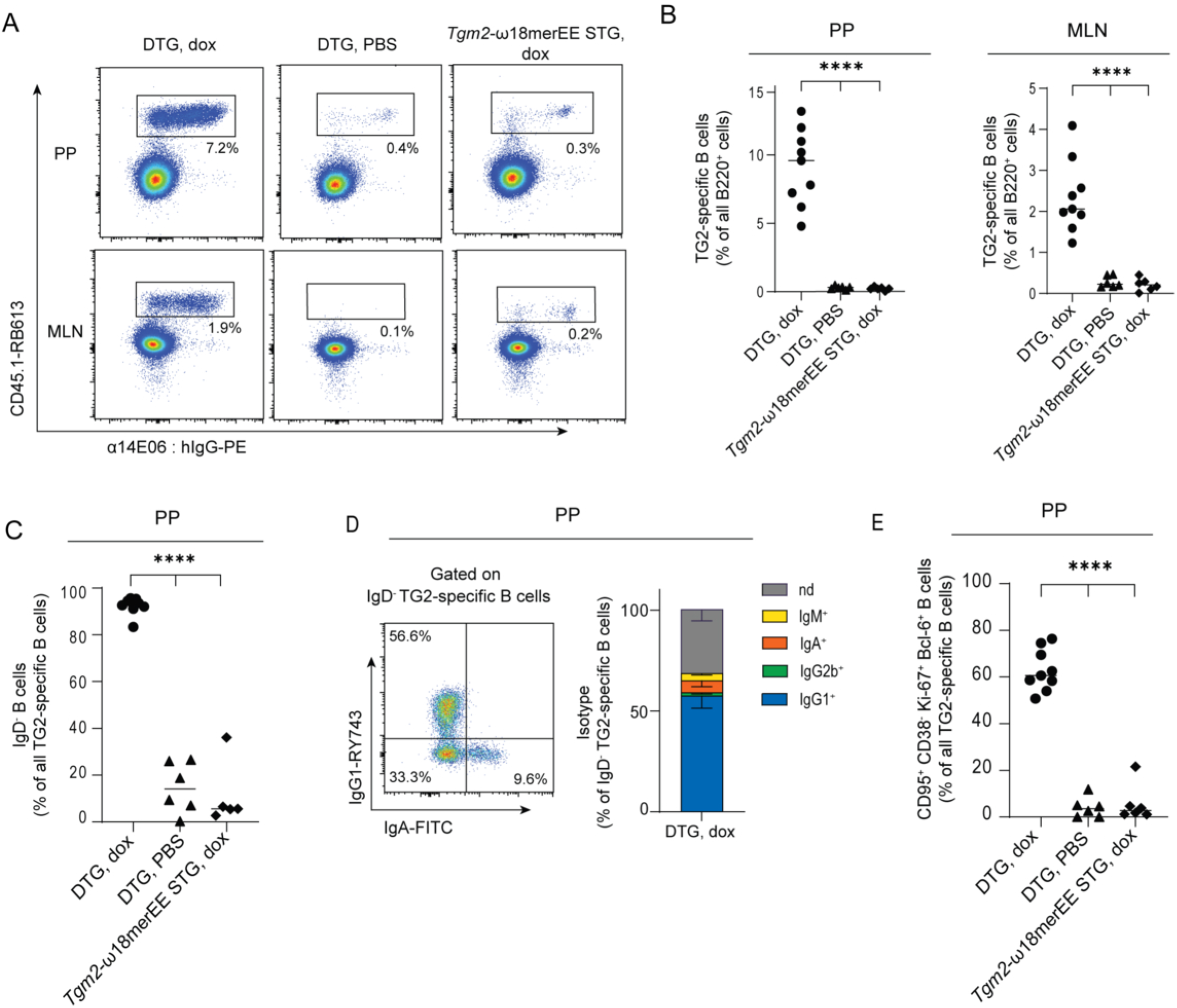
Strong expansion of TG2-specific B cells in response to IEC-derived TG2-ω18merEE fusion protein in Peyer’s patches. CD45.2 HLA-DQ2.5 DTG or CD45.2 HLA-DQ2.5 *Tgm2*-ω18merEE STG mice received naive gluten-specific CD4^+^ T cells, isolated from HLA-DQ2.5 TCR-glia-ω2 mice, by adoptive transfer and were subsequently immunized i.g. with DGP (500 µg) and CT (10 µg). Naive CD45.1 TG2-specific B cells, isolated from CD45.1 *Tgm2*^-/-^ HLA-DQ2.5 14E06 KI mice, were then transferred, followed by i.p. administration of dox (50 mg/kg) or PBS, as outlined in Figure 3A. Mice were euthanized 7 days after dox administration. **(A)** Representative plots of TG2-specific CD45.1^+^ B cells in Peyer’s patches (PPs; top) and mesenteric lymph nodes (MLNs; bottom). TG2-specific B cells are recognized by an anti-14E06 antibody targeting the 14E06 KI BCR. **(B)** Graphical representation of transferred TG2-specific B cells as a percentage of all B220^+^ B cells in PPs and MLNs. Data are pooled from four independent experiments. **(C)** Frequency of class switched (IgD^-^) TG2-specific B cells in PPs. Data are pooled from four independent experiments. **(D)** Immunoglobulin isotype distribution among IgD^-^ TG2-specific B cells in PPs. A representative scatter plot showing IgG versus IgA and a summary graph with pooled data shown as mean and SD from two independent experiments with a total of five mice included. N.d. not detected.. **(E)** Frequency of TG2-specific B cells in PPs with a germinal center (GC) phenotype, identified as CD95⁺ CD38⁻ Ki-67⁺ Bcl-6⁺. (**A-E**) Data are shown with median. Ns: not significant, *p < 0.05, **p < 0.01, ***p < 0.001, and ****p < 0.0001 as determined by one-way ANOVA. **Figure Supplement 2.** Flow data on activation and differentiation status of TG2-specific B cells in MLN, corresponding to Figure 4C,D,E. **Figure Supplement 3.** Gating strategy.

Transferred transgenic gluten-specific CD4^+^ T cells were identified by staining for the transgenic human TCR β chain Vβ3. As all mice were orally immunized with DGP and CT to prime transferred gluten-specific CD4^+^ T cells before IEC expression of TG2-ω18merEE was induced, these cells displayed an activated phenotype (CD62L^-^ CD44^+^) in both PPs and MLNs in all groups (Figure Supplement 2D). The activation of gluten-specific CD4^+^ T cells, as well as of TG2-specific B cells, was restricted to the gut as no B-cell and T-cell expansions were seen in peripheral lymph nodes (Figure Supplement 2E). While the frequency of gluten-specific CD4^+^ T cells in MLNs did not differ between mice with or without induced IEC expression of TG2-ω18merEE, their frequency was significantly increased in PPs of mice in which IEC expression of the fusion protein had been induced, that is, the group in which TG2-specific B cells showed robust expansion in PPs (Figure 5A). In these mice, the majority of the gluten-specific T cells had acquired a T follicular helper (T_FH_) phenotype (CD4^+^ CXCR5^+^ PD1^hi^ and Bcl-6^+^) (Figure 5B,C), strongly suggesting collaboration of these cells with TG2-specific B cells in PPs. In contrast, the frequency of gluten-specific CD4^+^ T cells with a T_FH_ phenotype was lower in MLNs (Figure Supplement 2F,G). Taken together these findings identify PPs as a dominant inductive site for the anti-TG2 response and strongly suggest that uptake and presentation of luminal, IEC-derived TG2-gluten antigen underlies the initiation of anti-TG2 autoimmunity.

**Figure 5.**
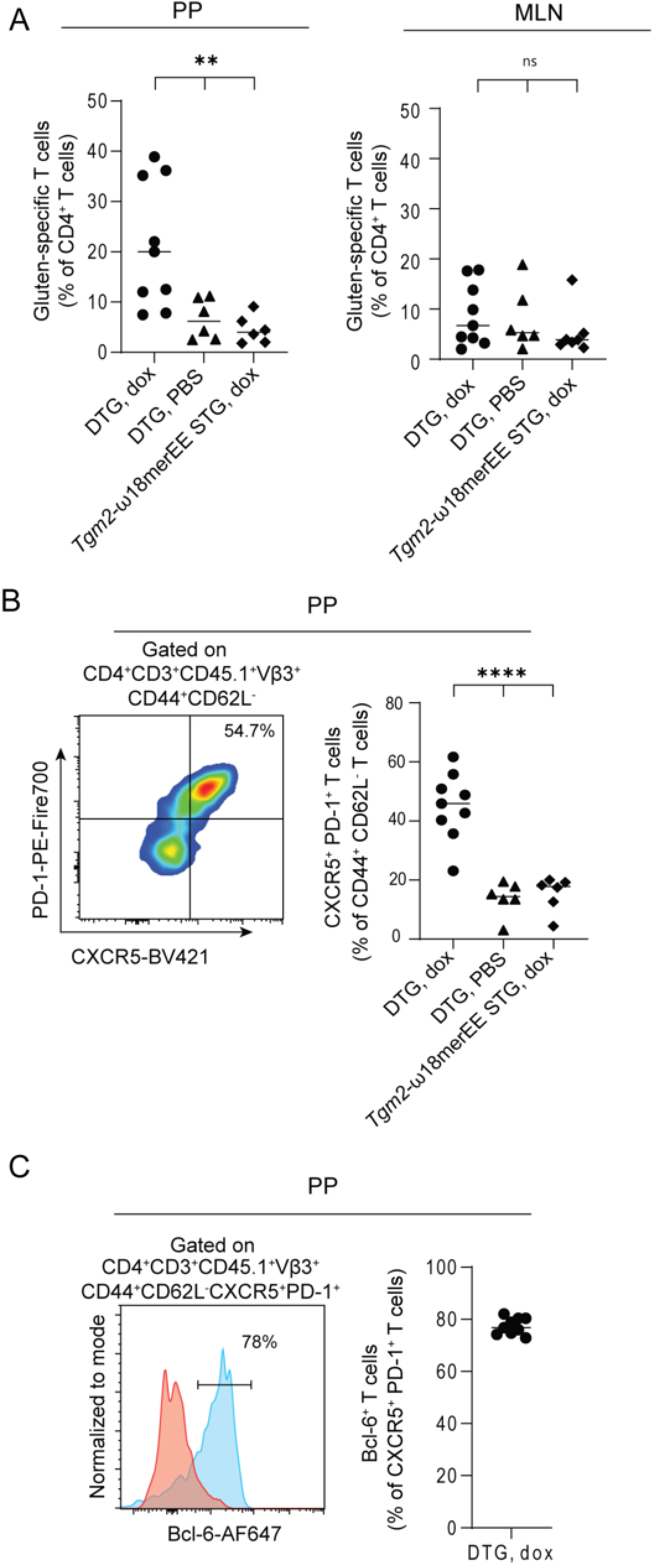
Gluten-specific T cells expand and adopt a T_FH_-phenotype upon expression of TG2-ω18merEE following transfer of TG2-specific B cells. Recipient mice received adoptively transferred gluten-specific CD4⁺ T cells, followed by immunization with DGP (500 μg) and CT (10 μg), adoptive transfer of TG2-specific B cells, and subsequent treatment with dox (50 mg/kg) or PBS. Mice were euthanized 7 days after dox administration. **(A)** Graphical representation of presence of gluten-specific TCR-glia-ω2 T cells as a percentage of all CD4^+^ T cells in PPs and MLNs. **(B)** Representative plot (left) and graphical summary (right) of the percentage of T cells with a T_FH_-like phenotype (CXCR5^+^, PD-1^+^) among the CD44^+^ CD62L^-^ gluten-specific CD4^+^ T cells in PPs. **(C)** Representative histogram plot (left) and graphical summary (right) showing the percentage of Bcl-6⁺ cells among gluten-specific T follicular helper (T_FH)_ cells. Bcl-6 expression within CXCR5^+^PD-1^+^-gated T_FH_ cells (blue) relative to the CXCR5^-^PD-1^-^ population (red). (**A-C**). Data are shown with median and are pooled from four independent experiments. Ns: not significant, *p < 0.05, **p < 0.01, ***p < 0.001, and ****p < 0.0001 as determined by one-way ANOVA. **Figure Supplement 2.** Supplementary data on activation and differentiation status of TG2-specific B cells and gluten-specific CD4^+^ T cells in gut-draining and peripheral lymph nodes. **Figure Supplement 3.** Gating strategy.

## DISCUSSION

We here present a mouse model in which anti-TG2 autoimmunity is driven by expression of a TG2-DGP fusion protein that is selectively induced in IECs. Our findings provide mechanistic support for a model in which TG2 derived from shed IECs forms transient complexes with gluten in the gut lumen and drives the collaboration between TG2-specific B cells and gluten-specific T cells in GALT (Iversen et al., 2020).

The previous observation that catalytically active TG2 is present in the gut lumen of mice relied on the addition of protease inhibitors to the analyzed gut lavage fluids (Meling et al., 2024). It remained uncertain from these observations whether TG2 released from shed IECs would survive in the gut lumen to reach immune inductive sites as protease inhibitors had to be added to lavage fluids in order to detect the TG2 protein. The experiments presented here prove that indeed it does. The BCR of the TG2-specific B cells recognizes a conformational epitope of TG2 (du Pre et al., 2020), and the only source of the induced TG2-ω18merEE fusion protein in the model system is IECs. The fusion protein’s ability to activate B cells implies that TG2 produced in the cytosol of IECs reaches immune inductive sites in an intact condition.

The striking expansion of transferred TG2-specific B cells observed in PPs of dox-treated DTG mice suggests that priming of TG2-specific B-cells occurs in PPs and possibly also in isolated lymphoid structures that are part of the inductive GALT. This is supported by the anatomical proximity of GALT to IECs shed into the gut lumen. Luminal antigens are rapidly sampled by specialized M cells in the follicle-associated epithelium of PPs that transport them directly to immune cells, including B cells (Komban et al., 2019), in the subepithelial dome. This short transit time ensures preservation of conformational epitopes. In active celiac disease small lymphoid aggregates associated with crypt regions, where B cells and T cells co-localize, may potentially also represent such immune inductive sites (FitzPatrick et al., 2025).

The formation of anti-TG2 antibodies in celiac disease is dependent on exposure to dietary gluten. When celiac disease patients commence a gluten-free diet, serum anti-TG2 IgA drops rapidly to levels of healthy subjects within weeks to months (Sulkanen et al., 1998) and TG2-specific plasma cells in the lamina propria of the disease lesion diminish to almost zero within months (Di Niro et al., 2016). The observation that gluten, a foreign antigen, controls autoantibody production laid the foundation for the model where gluten-specific CD4^+^ T cells provide T-cell help to autoreactive B cells through linkage of the self-antigen TG2 with a gluten T-cell epitope (Sollid et al., 1997). When gluten is removed from the diet, T-cell help for TG2-specific B cells will diminish and antibody titers drop. We have previously demonstrated that there is no B-cell tolerance to TG2 and that the formation of autoantibodies to TG2 is regulated at the level of T-cell help (du Pre et al., 2020). Our current findings that the development of anti-TG2 autoimmunity in the inducible TG2-DGP fusion protein model depends on the presence of activated gluten-specific T cells extend this notion and indicate that having an inflammatory gluten-specific CD4^+^ T-cell response is sufficient to induce the production of anti-TG2 autoantibodies when TG2-gluten complexes are present in the gut lumen. The finding in the mouse model that naive TG2-specific B cells readily become antibody producing cells in the lamina propria when provided T-cell help, fits with observations in celiac disease patients which suggest that the TG2 specific plasma cells are predominantly recruited from the naive B cell pool (Lindeman et al., 2024).

A requirement for the formation of luminal TG2-gluten complexes that can activate TG2-specific B cells is a sufficient amount of TG2 derived from shed IECs, together with a high concentration of gluten peptides. The availability of TG2 in the gut lumen will largely be determined by the turnover rate of the epithelial layer. In the human small intestine, the transit time from cell generation at the bottom of the crypts to cell expulsion at the villus apex is 3-5 days (Lipkin et al., 1963; Macdonald et al., 1964), and in patients with active celiac disease there is a three-fold increase in proliferating cells per crypt (Wright et al., 1973) and the IEC-turnover is shortened to 24 h (Savidge et al., 1995). Moreover, in the apical epithelial compartment from which IECs are shed into the lumen, the TG2 protein expression per cell is doubled in untreated celiac disease (Amundsen et al., 2023). Consistent with this, gene expression levels of TG2 were increased in shed IECs compared to tissue-resident cells in mice (Bahar Halpern et al., 2023). Furthermore, TG2 expression was shown to be increased following gluten challenge in human organoids (Dotsenko et al., 2024). Hence, the condition in active celiac disease is poised for generation of TG2-gluten enzyme-substrate complexes that can reach GALT and drive the generation of TG2-specific plasmablasts and gluten-specific CD4^+^ T cells that eventually home to the lamina propria of the celiac lesion as immune effector cells.

Immune reactivity to IEC-specific antigens has been studied previously. Vezys and coworkers made mice expressing cytosolic OVA selectively in IECs in a non-secreted form (Vezys et al., 2000). In these mice, transfer of OVA-specific CD8^+^ T cells led to expansion of these T cells in PPs and MLNs with subsequent migration of T cells to the epithelium, leading the authors to conclude that the T cells were activated by ovalbumin from shed IECs in PPs and MLNs. Despite T-cell expansion, no tissue damage was observed, suggesting that the T cells adopted a tolerogenic state (Vezys et al., 2000). However, upon inflammation, caused by infection with vesicular stomatitis virus with or without ovalbumin expression, or by other inflammatory stimuli (Vezys and Lefrancois, 2002; Vezys et al., 2000), overt T-cell mediated destruction of IECs ensued. It was speculated that a mechanism where dendritic cells in the intestinal mucosa take up apoptotic IECs and migrate to T cell areas of MLNs (Huang et al., 2000) could be responsible for the systemic tolerance. Similarly, Cummings et al. observed uptake of apoptotic IECs by dendritic cells and two macrophage subsets in the lamina propria with the migratory dendritic cells inducing CD4^+^ T regulatory cells in MLNs in mice (Cummings et al., 2016). It is conceivable that this mechanism of uptake of apoptotic IECs expressing the TG2-DGP fusion protein also occurs in our mice, yet this pathway can hardly explain the presentation of conformationally intact TG2 antigen to B cells. Importantly, the findings of Vezys and coworkers support the notion that intracellular antigen expressed by IECs can indeed instigate a response in PPs.

Gut luminal TG2 is a potential therapeutic target in celiac disease. The TG2 at this location is unlikely to have a physiological function and designing a drug to act locally in the gut without systemic absorption would minimize the likelihood of unwanted side effects of the treatment. This is certainly an advantage; however, because TG2 is continuously released from a large number of shed IECs, sustained quenching is difficult to achieve. In this setting it is notable that TAK227/ZED1227, an irreversible TG2 inhibitor with little systemic absorption (Buchold et al., 2022), was demonstrated to reduce inflammation and tissue remodeling in celiac disease patients who received the drug perorally 30 min ahead of eating a gluten containing muesli bar once daily for 6 weeks (Schuppan et al., 2021).

This experimental work demonstrates that TG2 selectively expressed in IECs is implicated in the generation of an intestinal B-cell response to TG2 when T-cell help is provided. The expressed TG2 fusion protein antigen is made to mimic the TG2-gluten peptide complex by covalent attachment of the deamidated gluten T-cell epitope to the C-terminus of TG2, which underscores that gluten-specific T cells can give the required T-cell help for the activation of TG2-specific B cells. The model generated will be useful to dissect the T-cell and B-cell crosstalk involved in the pathogenesis of celiac disease and to test therapies aiming to interfere with the adaptive immune responses causing celiac disease.

## MATERIALS AND METHODS

### Mice

Rosa26 *Tgm2*-ω18merEE H2B-eGFP mice were generated by Ozgene Pty Ltd. Briefly, a gene construct was generated composed of the cDNA sequences of murine TG2 (ENSMUST00000103122.9) with the retention of intron 7 fused to the 18 residue ω-gliadin peptide QPEQPFPQPEQPFPWQPQ via a 3x glycine linker in reverse orientation, P2A from porcine teschovirus, the H2B-eGFP reporter gene (Tumbar et al., 2004), pA terminal sequence from bovine growth hormone, and a neomycin resistance cassette flanked by loxP sites (Figure 1A). The construct was electroporated into C57Bl/6 embryonic stem cells. Neomycin resistant embryonic stem cells were injected into goGermline blastocysts (Koentgen et al., 2016). The generated mice were crossed to flippase-expressing B6J-*Gt(ROSA)26Sor*^tm1(UbiC-FLPe)Ozg/Ozg^ mice to delete the neomycin cassette. The resulting *Tgm2*-ω18merEE transgenic mice were crossed with B6;SJL-Tg(Vil1-rtTA*M2)5Stang/J mice (IMSR_JAX:031285) (Chen et al., 2014) to generate *Tgm2*-ω18merEE IEC-rtTA DTG mice. For adoptive transfer experiments, DTG mice were crossed to HLA-DQ2.5 KI mice (Dewan et al., 2021). TCR-glia-ω2 transgenic mice (Lindstad et al., 2021) and 14E06 KI mice (du Pre et al., 2020) crossed to HLA-DQ2.5 mice are described elsewhere. To avoid exposure of TG2-specific B cells to TG2 antigen released during tissue processing and cell isolation, HLA-DQ2.5 14E06 KI mice crossed to *Tgm2*^-/-^ mice (De Laurenzi and Melino, 2001) were used as B-cell donors in all experiments. For certain experiments, HLA-DQ2.5 14E06 KI were crossed to CD45.1 mice (B6.SJL-Ptprca Pepcb /BoyCrl, Charles River Laboratories), when indicated. Recipient mice of both sexes were used, with donor and recipient sex matched except that female donor cells were permitted in male recipients. All strains were bred on a C57Bl/6 background, maintained on irradiated gluten-free chow (D11112201i, Research Diets), and kept under specific-pathogen free conditions on a 12 h light/dark cycle at the Department of Comparative Medicine, Oslo University Hospital, Rikshospitalet, Oslo, Norway. All experiments were approved by the Norwegian Food Safety Authority (Mattilsynet) and conducted in accordance with Directive 2010/63/EU.

### Immunohistochemistry

DTG mice received dox (50 mg/kg in PBS) by i.p. injection and were euthanized after 24 h. Approximately 3 cm segments of small intestine were flushed with PBS, fixed in 4% paraformaldehyde and 10% sucrose for 1 h on ice, and transferred to 30% sucrose for 16–48 h at 4 °C. Samples were embedded in Tissue-Tek OCT compound, snap-frozen in liquid nitrogen, and sectioned at 10 μm. Tissue sections were blocked with 1.25% IgG-free BSA (Jackson ImmunoResearch) in PBS, and stained with 4′,6-diamidino-2-phenylindole (DAPI,62248, Invitrogen), anti-ω18merEE (PD93), rabbit anti-mouse TG2 (Pacific Immunology), donkey anti-rabbit IgG-Cy3 (711-165-152, Jackson ImmunoResearch), and goat anti-human IgG SA647 (A-214445, ThermoFisher) at room temperature. Slides were mounted with ProLong™ Diamond Antifade Mountant (P36961, Invitrogen). Images were acquired at room temperature on a Zeiss LSM 980 confocal microscope using 20x NA 0.8 DIC II and 63x NA 1.4 oil DIC III (Plan-Apochromat) objectives and ZEN microscopy software. Image brightness and contrast were adjusted using Fiji/ImageJ software.

### Intestinal epithelial cell lysate preparation

DTG or *Tgm*2-ω18merEE or IEC-rtTA STG mice received dox (2 mg/ml, D3447, Sigma-Aldrich) in drinking water supplemented with 2 % (w/v) sucrose to mask the bitter taste, for three days followed by one day of normal drinking water before euthanasia. The small intestine was carefully removed, then extensively washed and small pieces collected in Ca^2+^- and Mg^2+^-free (CMF) HBSS (14180-046, Gibco) supplemented with HEPES (10 mM, H0887, Sigma-Aldrich). IECs were isolated by incubation in CMF HBSS supplemented with HEPES (10 mM), EDTA (5 mM, 15575038, Invitrogen), 0, 5% (v/v) FCS (F7524, Sigma-Aldrich), and dithiothreitol (DTT, 1 mM, D9779, Sigma-Aldrich) for 20 min at 37 °C under continuous rotation. Cells were collected and resuspended in CMF HBSS supplemented with EDTA (1 mM) before lysis by three freeze/thaw cycles. Lysed samples were centrifuged at 16,200*g* for 10 min at 4 °C and supernatants stored at -70 °C.

### Isolation of T cells and B cells

Single-cell suspensions were prepared by carefully grinding lymph nodes and spleens and filtering over a 70 µM mesh. Erythrocytes were lysed by treatment with ammonium chloride potassium lysis buffer. B cells were isolated from *Tgm2*^-/-^ HLA-DQ2.5 14E06 KI mouse spleens using Dynabeads Mouse CD43 kit (11422D, Invitrogen). CD4^+^ T cells were isolated from lymph nodes and spleens of TCR-glia-ω2 mice using the Easysep Mouse CD4^+^ T cell isolation kit (19852, StemCell Technologies).

### *In vitro* B-cell and T-cell proliferation assay

Isolated TG2-specific B cells were labeled with cell trace violet proliferation-tracking dye (CTV, C34557, Invitrogen) according to the manufacturer’s instruction and pulsed with IEC-lysate for 5 mins on ice. Isolated gluten-specific CD4^+^ Vß3^+^ T cells were labeled with CFSE (65-0850-85, eBioScience), according to the manufacturer’s instruction. B cells were then co-cultured at 2 x 10^5^ B220^+^ cells/well with 2.5 x 10^4^ CD4^+^ cells/well in RPMI 1640 (R2405, Sigma-Aldrich) supplemented with 10% (v/v) FCS, 1% (v/v) penicillin/streptomycin (P4333, Sigma-Aldrich), 2-ME (100 µM, M3148, Sigma-Aldrich), sodium pyruvate (1 mM, 11360-070, Gibco), nonessential amino acids (0.1 mM, M7145, Sigma-Aldrich), and HEPES (10 mM) at 37 °C and 5% CO_2_. After 3 days, proliferation levels were determined by flow cytometry.

### Immunoprecipitation

To extract TG2-ω18merEE fusion protein from the gut lumen, small-intestinal lavage fluids were collected by clamping the small intestine at one end and injecting 1 ml PBS supplemented with soybean trypsin inhibitor (SBTI, 2 mg/ml, T9128, Sigma-Aldrich) and Pefabloc (4 mM, 11585916001, Roche) using a syringe with a smooth-tipped feeding needle (1044, AgnThos), then clamping the other end. Small intestines were incubated for 10 mins in a PBS-filled petri dish on ice before lavage fluid was collected and snap frozen in liquid N_2_. Protein G Dynabeads (25 mg/ml, 10003D, Invitrogen) were incubated with celiac patient-derived recombinant monoclonal hIgG1 anti-DGP antibody (0.17 mg/ml, UCD1002-1E03) in Tris-buffered saline (TBS, pH 7.4) with 0.02% Tween (P1379, Sigma-Aldrich) for 10 mins at room temperature under continuous agitation. Excess antibody was removed using magnetic separation. Lavage fluids were centrifuged at 11 600*g* for 10 mins and supernatants added to the beads and diluted to a total volume of 1 ml with TBS, before being incubated for 30 mins at room temperature under continuous agitation. TG2-ω18merEE fusion protein was eluted from the beads by 2x Laemmli buffer (20% glycerol, 4% SDS, 0.02% bromophenol blue, 125 mM Tris-HCl pH 6.8) supplemented with 2.5% (v/v) β-ME and boiled for 7 mins. Supernatants were collected for SDS-PAGE.

### Expression of anti-DGP human monoclonal antibodies

The hIgG1 UCD1002-1E03 anti-DGP recombinant monoclonal antibody derived from a human plasma cell (Steinsbo et al., 2014) was expressed in Expi293F cells (A14527, Thermofisher Scientific) as previously described (Smith et al., 2009). Briefly, Expi293F cells cultured in F17 medium (A13835-01, Gibco) were expanded to 3 x 10^6^ cells/ml in shaker flasks maintained under continuous agitation at 110 rpm at 37 °C with 8% CO_2_. Cells were transfected with 0.1 mg/ml polyethyleneimine (PEI, 23966, Polysciences), 0.25 µg/ml IgH plasmid, and 0.625 µg/ml IgL plasmid DNA incubated at room temperature for 15 mins before adding to the cell suspension. Expressing cells were cultured for 5 days, then harvested by centrifugation at 945 x *g* for 30 mins at 4°C, and the resulting supernatant was clarified by further centrifugation at 4000 x *g* for 20 mins at 4°C, and filtration with a 0.22 *μ*m polyethersulfone sterile filtration unit. The antibody was affinity purified using a 1ml HiTrap Protein G column (17-0404-01,GE Healthcare) on an Äkta Pure by applying the clarified filtrate to the column, then washing with 5 column volumes of 1 M NaCl, then 15 column volumes of 20 mM NaP (pH 7.5), then antibody was eluted with 100 mM glycine (pH 2.7) in 0.5ml aliquots. 30 µl 1 M Tris-HCl (pH 9) was added to each aliquot collection tube. 1.5 ml of purified protein was dialyzed against 3 L PBS in a 0.5 – 3 ml 20 kDa molecular weight cut-off (MWCO) Slide-A-Lyzer dialysis cassette (66003, Thermofisher Scientific) to a final protein concentration of 8.2 mg/ml, and stored in aliquots at -20°C.

Anti-gliadin antibody PD93 was modified from anti-gliadin antibody UCD1002-1E03 to increase its affinity using phage display (Heggelund et al., manuscript in preparation). The recombinant hIgG1 PD93 was expressed in Expi293F cells cultured in EXPI293 Expression Medium (13469756, Thermofisher Scientific) using shaker flasks maintained at 37 °C with 8% CO₂. Twenty-four hours prior to transfection, cells were diluted to a density of 1 × 10^6^ cells/mL. Transfection was performed by co-diluting 25 *μ*g each of IgH and IgL chain plasmid DNA with 0.1 mg/ml PEI in 5 mL of Expi293 Expression Medium. The mixture was incubated for 15 min at room temperature before addition to the cell suspension. Expressing cultures were maintained under continuous agitation at 110 rpm for 6 days at 37 °C with 8% CO₂.Cells were harvested by centrifugation at 3,000 × *g* for 20 min at 4 °C, and the resulting supernatant was clarified using a 0.22 *μ*m polyethersulfone sterile filtration unit. For affinity purification, 0.5 mL of CaptureSelect CH1-XL affinity matrix (1943462005, ThermoFisher) was added to the clarified filtrate and batch-incubated for 1.5 h at 4 °C on a tube roller. The resin was captured in a gravity flow column, washed with 10 column volumes (5 mL) of 20 mM sodium phosphate (pH 7.2), and eluted with 100 mM glycine (pH 2.7). Elution fractions were immediately neutralized by adding 1 M Tris-HCl (pH 8.0). The antibody was further purified by size exclusion chromatography using a Superdex 200 10/300 GL column (Cytiva) running at 0.75 ml/min in PBS. The purified protein was concentrated to 0.55 mg/ml using a 10 kDa molecular weight cut-off (MWCO) and stored in aliquots at -20°C.

### Western blotting

Immunoprecipitated proteins or IEC-lysates were separated by SDS-PAGE. IEC-lysates were diluted in Laemmli sample loading buffer (10% glycerol, 2% SDS, 0.005% bromophenol blue, 62.5 mM Tris-HCl pH 6.8, 2% β-ME, final concentrations) and boiled for 5 mins before all samples were loaded into a 4-20% Miniprotean TGX gel run in 1 x SDS running buffer (25 mM Tris, 192 mM Glycine, 0.1% SDS). A recombinant TG2-ω18merEE fusion protein expressed as previously described (Loberg et al., 2025), was loaded as a control. The gel was equilibrated for 15 mins in Towbin transfer buffer (25mM Trizma base, 192mM Glycine, 20% Methanol, pH 8.3), before semi-dry transfer (Biorad) to a nitrocellulose membrane (10600020, Cytiva). The membrane was blocked in 5% skim milk in TBST-buffer (50 mM Tris pH 7.6, 150 mM NaCl, 0.1% (v/v) Tween) for 1 h at room temperature, followed by incubation with primary antibodies in 1.25% skim milk in TBST-buffer at 4 °C overnight. Primary antibodies were custom rabbit–anti-mouse TG2 polyclonal antibody (1:40,000, Pacific Immunology) for detecting protein from cell lysate and immunoprecipitated lavage fluid, and rabbit–anti-β-actin (D6A8) antibody (1:3000, 8457, Cell Signaling Technology) and hIgG1 anti-DGP antibody (1:2000, UCD1002-1E03) for detecting protein from cell lysate. After TBST-wash (3 x 10 min), the membrane was incubated with horseradish peroxidase-conjugated goat–anti-rabbit IgG (1:3000, 4010-05, Southern Biotech) and horseradish peroxidase-conjugated F(ab’)₂ fragment donkey-anti-human IgG (1:5000, 709-036-098, Jackson ImmunoResearch Laboratories Inc.) for 1 h at room temperature, followed by an additional 3 x 10-min wash in TBST-buffer. The membrane was developed with Super signal West Pico Plus (ThermoFisher Scientific) using a chemiluminescence reader (G:BOX Chemi XRQ, Syngene).

### Adoptive transfer and immunization

Isolated gluten-specific CD4^+^ T cells were adoptively transferred into HLA-DQ2.5 DTG mice in 200 µL PBS via injection in the lateral tail vein. The following day, recipient mice received ω18merEE peptides (100 µg or 500 µg, as indicated, QPEQPFPQPEQPFPWQPQ, Genscript) and CT (10 µg, *Vibrio cholerae*, 100B, List Biological Laboratories) in 200 µL PBS with 3% (w/v) NaHCO3 (Sigma-Aldrich) via i.g. administration. Three days later, isolated TG2-specific B cells were transferred by i.v. injection and after 2 h, the recipient mice were administered dox (50 mg/kg of body weight) via i.p. injection. Mice were euthanized 7 or 14 days (as indicated) after dox treatment.

### ELISA

Blood was collected from the saphenous vein and allowed to coagulate at room temperature for 1 h before centrifugation at 2000 x *g* for 15 mins at 4 °C. Serum was collected and stored at -20 °C. To sample antibodies in the intestinal lumen, small-intestinal lavage fluid was collected by clamping one end of the small intestine and injecting 1.5 ml PBS supplemented with SBTI (0.1 mg/ml), EDTA (50 mM), and Pefabloc (4 mM) using a syringe with a smooth-tipped feeding needle (1044, AgnThos), followed by clamping the other end. The intestines were then incubated for 5 mins in a PBS-filled petri-dish on ice before lavage fluid was collected and centrifuged at 840*g*, 4 °C for 10 mins. Pefabloc (400 µM) was added to the supernatants before centrifuging at 16,200 x *g* for 15 mins at 4°C. Pefabloc (400 µM) and 0.01% (v/v) sodium azide were added to supernatants and incubated for 15 mins on ice before adding 3% (v/v) FCS and centrifuging at 16,200 x *g* for 15 mins at 4 °C. Supernatants were stored at -70° C. For detection of anti-TG2 antibodies, 96-well Maxisorp ELISA plates (Nunc) were coated overnight at 4°C with 50 µL of a recombinant mouse TG2 (5 µg/ml) expressed in Sf^+^ express insect cells as previously described (du Pre et al., 2020).

Recombinant mouse TG2 was immobilized in an open conformation using the irreversible active site inhibitor DP3-3 (66.5 µg/ml, Ac-P(DON)LPF-NH_2_; Zedira). After washing, 50 µL of 5-fold serial dilutions of serum starting from 1:50 or of small intestinal lavage fluid starting from 1:2 in PBS supplemented with 0.01% (v/v) Tween and 0.5% (v/v) FCS was added to wells and plates were incubated for 2 h at room temperature. Plates were washed, then antibodies were detected by incubating with alkaline phosphatase (AP)-conjugated goat anti-mouse IgG (A2429, Sigma-Aldrich) or goat anti-mouse IgA (1040-04, Southern Biotech) for 1 h at room temperature, before washing and incubating with AP substrate (34047, Thermo Fisher Scientific). Absorbance was measured at 405 nm using a plate reader (Multiskan ascent, ThermoFisher Scientific). Area under the curve was determined as a function of the ratio of serum or lavage fluid to buffer and is measured as peaks above baseline 0, ignoring peaks less than 10% of the maximum peak, using the GraphPad Prism 10 software.

### ELISPOT

Small intestines, with PPs removed, were washed in CMF HBSS before two rounds of incubation in CMF HBSS supplemented with HEPES (10 mM), EDTA (5 mM), 5% (v/v) FCS, and DTT (1 mM) for 15 mins at 37 °C under continuous rotation. After washing with CMF HBSS supplemented with HEPES (10 mM), to remove residual EDTA, lamina propria lymphocytes were isolated by digesting the tissue using 0.5 mg/ml Collagenase VIII (C2139, Sigma-Aldrich) in HBSS supplemented with HEPES (10 mM) and 5% (v/v) FCS and running program 37C_m_LPDK_1 on a gentleMACS™ Octo Dissociator with heaters (Miltenyi Biotec). Multiscreen HPS membrane plates (Millipore) were activated with 20 µL 35% (v/v) ethanol for 1 min at room temperature before washing 3x with 200 µL PBS. Plates were coated with 100 µL 5 µg/ml NeutrAvidin (31000, Thermo Fisher Scientific) or anti-mouse IgA (556960, BD Pharmingen) at 4 °C overnight. After washing and blocking, NeutrAvidin-coated plates were incubated for 1 h at room temperature with 100 µL biotinylated human TG2. Biotinylated recombinant human TG2 was produced in Sf^+^ insect cells, as previously described (Das et al., 2024). Isolated lamina propria lymphocytes were added to the plates in three-fold serial dilutions, in RPMI 1640 supplemented with 10% (v/v) FCS, 1% (v/v) penicillin/streptomycin, and β-ME (100 µM), starting from 2 x 10^5^ cells when detecting for anti-TG2 plasma cells and 2 x 10^4^ when detecting for all IgA-producing plasma cells. Plates were incubated for 3 h at 37 °C and 5% CO_2_ before washing with PBS with 0.01% (v/v) Tween and incubating at 4°C overnight with anti-mouse IgA-AP (1:1000). Plates were extensively washed with PBS-T (0.01% (v/v) Tween), PBS, and dH_2_O before detecting spots with NBT-5-bromo-4-chloro-3-indolyl phosphate substrate (170-6432, Bio-Rad). Plates were rinsed thoroughly in dH_2_O and dried overnight. Plates were scanned using a Cellular Technologies Limited (CTL) Immunospot Analyzer, and spots were quantified using the CTL Immunospot 5.1 software. Wells incubated in medium served as negative controls.

### Flow cytometry

Single-cell suspensions of PPs were made by grinding and filtering over a 70 µm mesh into PBS with 2% (v/v) FCS and EDTA (2 mM). Single-cell suspensions of lymph nodes were prepared by roughly cutting in PBS with 2% (v/v) FCS with surgical scissors then carefully grinding them over a 70 µM mesh. MLNs, peripheral LNs (axial and brachial) and PPs were incubated in PBS with 2% (v/v) FCS and EDTA (2 mM) and the Fc receptor blocked using an anti-FcII/II antibody (anti-mouse CD16/32, clone 93; BioLegend). Dead cells were identified using Zombie NIR Fixable Viability Kit (11863, Sony). Cells were fixed and permeabilized using eBioscience™ Foxp3 / Transcription Factor Staining Buffer set (Invitrogen), according to the manufacturer’s instruction. Antibodies used for staining were B220 – allophycocyanin (APC), clone RA3-6B2, BioLegend, B220 – Brilliant Ultra Violet (BUV) 496 (clone RA3-6B2, BD), Bcl-6 – Alexa Fluor (AF) 647 (clone K112-91, BD), CD3 – BUV 395 (clone 145-2C11, BD), CD4 – APC-Cy7 (clone GK1.5, BioLegend), CD4 – APC-Fire810 (clone GK1.5, BioLegend), CD44 – Brilliant Violet (BV) 605 (clone IM7, BioLegend), CD45.1– RealBlue (RB) 613 (clone A20, BD), CD45.2 – APC-Cy7 (clone 104, BioLegend), CD62L – SparkRed718 (clone MEL-14, BioLegend), CD95 (Fas) – PE-Cy7 (clone Jo2, BD), CD185 (CXCR5) – BV421 (clone L138D7, BioLegend), CD279 (PD-1) PE-Fire700 (clone 29F.1A12; Sony Biotechnology), IgA – FITC (clone mA-6E1, eBioscience), IgD – BV786 (clone 11-26c.2a, BD), IgG1 – Real Yellow (RY) 743 (clone A85-1, BD), IgG2b – RB545 (clone R12-3, BD), IgM – BUV805 (clone II/41, BD), Ki-67 – BV480 (clone B56, BD), and human TCR Vβ3/R – PE (clone JOVI-3, Ancell). Unconjugated hIgG1 anti-14E06 (used at 0,58 µg/ml) was expressed as previously described (Høydahl et al., 2016). Anti-14E06 was detected with anti-human IgG1-BV711 (clone M1310G05, BioLegend). Cells were analyzed on Attune Nxt Flow Cytometer or Sony ID 7000 Spectral Cell Analyser, followed by analysis using FlowJo software (BD).

## DATA AVAILABILITY

All data are available in the main text or supplementary materials.

## ACKNOWLEDGEMENTS

We thank Marie K. Johannesen for production of recombinant TG2 proteins. We are grateful to the Department of Comparative Medicine, Oslo University Hospital, for animal husbandry, and to the OUH HSØ Advanced Light Microscopy Core Facility for microscopy services.

## AUTHOR CONTRIBUTIONS

R.I. Løberg designed and performed experiments, analyzed data, and wrote the original draft. H.A. Abdi-Dezfuli designed and performed experiments and analyzed data. L. Kleppa performed experiments and contributed to manuscript revision. A. Dewan performed mouse experiments and contributed to manuscript revision. M.T. Meling performed experiments and contributed to manuscript revision. L.M. Sollid conceptualized and supervised the study, acquired funding, and contributed to manuscript revision. M.F. du Pré conceptualized and supervised the study, designed and performed experiments, acquired funding, and contributed to manuscript revision.

The authors declare that no competing interests exist

## ABBREVIATIONS

BCR: B-cell receptor
CT: cholera toxin
DGP: deamidated gluten peptide
Dox: doxycycline
DTG: double transgenic
GALT: Gut-associated lymphoid tissue
IEC: intestinal epithelial cell
i.g.: intragastric
KI: knock-in
MLN: mesenteric lymph node
PP: Peyer’s patch
rtTA: reverse tetracycline-controlled transactivator
STG: single transgenic
T_FH_: T follicular helper cell
TCR: T-cell receptor
TG2: transglutaminase 2
TG2-DGP: a fusion protein of TG2 and DGP
TRE: tetracycline responsive element

## Supplementary Figure legends

**Figure Supplement 1.**

**Gating strategy for i*n vitro* T-cell and B-cell proliferation assay, supporting Fig 1 E.** Gating of proliferated cells at day 3 of a co-culture assay of CTV-labeled TG2-specific B cells and CFSE-labeled gluten-specific CD4^+^ T cells in response to IEC-lysates of dox-treated DTG mice. Cells are first gated on lymphocytes (FSC-A x SSC-A scatter) and single cells (FSC-A x FSC-H scatter). T cells are first identified as B220^-^ and CTV^-^ before gating on CD4 and the transgenic human Vβ3 TCR. B cells are identified as CD4^-^ and CFSE^-^ before gating on B220^+^.

**Figure Supplement 2.**

**Supplementary data on transferred antigen-specific B cells and T cells, supporting Figures 4 and 5.** Recipient DTG mice received gluten-specific T cells and by intragastric administration of DGP (500 µg) and CT (10 µg) 24 hours after cell transfer. After 3 days, TG2-specific B cells were transferred and recipient DTG mice received dox (50 mg/kg i.p.), as illustrated in Fig 3 A. **(A)** Percentage of class-switched (IgD^-^) TG2-specific B cells in the MLNs. Data are pooled from four independent experiments. **(B)** Representative scatter plot (left) showing the IgG and IgA isotype distribution among IgD^-^ TG2-specific B cells, with the accompanying graph (right) summarizing the distribution of all immunoglobulin isotypes. Data shown as mean and SD are pooled from two independent experiments with a total of five mice included. N.d. not detected. **(C)** Frequency of TG2-specific B cells in the MLNs exhibiting a germinal center phenotype (CD95⁺ CD38^-^ Ki-67⁺ Bcl-6⁺). Data are pooled from four independent experiments. **(D)** Graphical representation of the percentage of activated (CD62L^-^, CD44^+^) gluten-specific TCR-glia-ω2 T cells in Peyer’s patches (PPs) and MLNs. Data are pooled from four independent experiments **(E)** Frequency of TG2-specific B cells (left), gated on B220^+^, and gluten-specific T cells (right), gated on CD4^+^ in peripheral (non gut-draining) lymph nodes. Data are pooled from two independent experiments. **(F)** Representative plot (left) showing gluten-specific T cells with a T_FH_-like phenotype (CXCR5^+^PD-1^+^), gated on activated T cells (CD44^+^CD62L^-^), with a graph (right) summarizing the percentage of gluten-specific T cells with this phenotype. Data are pooled from four independent experiments. **(G)** Representative histogram (left) showing Bcl-6 expression within of CXCR5^+^PD-1^+^-gated T_FH_ cells in MLNs (blue) compared to the CXCR5^-^ PD-1^-^ population (red). Graph shows quantification of the percentage of Bcl-6^+^ cells of four independent experiments. Ns: not significant, *p < 0.05, **p < 0.01, ***p < 0.001, and ****p < 0.0001 as determined by one-way ANOVA.

**Figure Supplement 3.**

**Gating strategy of adoptively transferred T and B cells, supporting Figures 4, 5 and S2.** Gluten-specific T cells, isolated from TCR-glia-ω2 mice, were transferred to CD45.2 HLA-DQ2.5 DTG mice and 24 hours later primed by intragastric administration of DGP (500 µg) and CT (10 µg). After 3 days, CD45.1^+^ TG2-specific B cells isolated from *Tgm2*^-/-^ HLA-DQ2.5 14E06 KI mice were transferred and DTG mice were treated with dox (50 mg/kg) via i.p. injections 2 hours after transfer. These plots are from PPs, and the gating strategies shown are representative of those used for both PPs and MLNs. **(A)** Viable cells were recollected in PPs and MLNs by gating on lymphocytes (FSC-A x SSC-A scatter), single cells (FSC-A x FSC-H scatter), and live cells (Zombie-NIR x FSC-A). **(B)** Adoptively transferred B cells were recollected in PPs and MLNs by gating on viable lymphocytes (A) and B cells (B220^+^). TG2-specific B cells were identified by a human IgG α14E06 followed by an anti-human IgG-PE. The isotype of class-switched cells w identified as IgD^-^IgG1^+^IgA^+^IgM^+^. Germinal center cells were determined as CD38^-^CD95^+^Ki-67^+^Bcl-6^+^ **(C)** Adoptively transferred T cells were identified in PPs and MLNs by gating on live lymphocytes (A) and CD3^+^CD4^+^ T cells. Gluten-specific T cells were identified by the human transgenic TCR Vβ3, activated T cells as CD44^+^ and CD62L^-^, and the T_FH_ phenotype as CXCR5^+^ and PD-1^+^. Cells with a T_FH_ phenotype (CXCR5^+^PD-1^+^) were determined to be Bcl-6^+^ (blue) by using CXCR5^-^PD-1^-^ as a negative control (red).

